# Butyrate Protects against SARS-CoV-2-induced Tissue Damage in Golden Hamsters

**DOI:** 10.1101/2023.07.27.550811

**Authors:** Huan Yu, Lunzhi Yuan, Zhigang Yan, Ming Zhou, Jianghui Ye, Kun Wu, Wenjia Chen, Rirong Chen, Ningshao Xia, Yi Guan, Huachen Zhu

## Abstract

Butyrate, produced by gut microbe during dietary fiber fermentation, plays anti-inflammatory and antioxidant effects in chronic inflammation diseases, yet it remains to be explored whether butyrate has protective effects against viral infections. Here, we demonstrated that butyrate alleviated tissue injury in severe acute respiratory syndrome coronavirus 2 (SARS-CoV-2)-infected golden hamsters with supplementation of butyrate before and during the infection. Butyrate-treated hamsters showed augmentation of type I interferon (IFN) response and activation of endothelial cells without exaggerated inflammation. In addition, butyrate regulated redox homeostasis by enhancing the activity of superoxide dismutase (SOD) to inhibit excessive apoptotic cell death. Therefore, butyrate exhibited an effective prevention against SARS-CoV-2 by upregulating antiviral immune responses and promoting cell survival.

**IMPORTANCE:** Since SARS-CoV-2 has caused severe disease characterized by acute respiratory distress syndrome (ARDS) in humans, it is essential to develop therapeutics based on relieving such severe clinical symptoms. Current therapy strategies mainly focus on individuals who have COVID-19, however, there is still a strong need for prevention and treatment of SARS-CoV-2 infection. This study showed that butyrate, a bacterial metabolite, improved the response of SARS-CoV-2-infected hamsters by reducing immunopathology caused by impaired antiviral defenses and inhibiting excessive apoptosis through reduction in oxidative stress.

## INTRODUCTION

Short-chain fatty acids (SCFAs) are the most abundant metabolites mainly produced by the gut microbiota in colon via fermentation of dietary fiber (1, 2). Among SCFAs, butyrate is a primary energy source for colonocytes and a well-known anti-inflammatory mediator, which can be activated by binding to G protein-coupled receptors (GPRs), mainly GPR41 and GPR43 (3), or inhibit the activity of histone deacetylase (HDAC) (4). Butyrate can not only regulate mucosal barrier function and mucosal immunity, but also mediate the communication between colonic microbiota and other organs such as brain, lung and liver (5–8).

Lower respiratory infections are reported the 4th leading cause of death with 2.6 million global deaths in 2019 (9). Since December 2019, severe acute respiratory syndrome coronavirus 2 (SARS-CoV-2) has caused more than 6.9 million deaths of coronavirus disease 2019 (COVID-19) by July 2023 (10). SARS-CoV-2 can replicate both in the upper and lower respiratory tract (11, 12). SARS-CoV-2 infection, alongside individual susceptibility and host immunity, can even progress to severe and life-threatening pneumonia, which is responsible for increased morbidity and mortality in COVID-19 (13). Patients with either acute COVID-19 or Post-acute COVID Syndrome (PACS) were reported to have gastrointestinal symptoms such as abdominal pain, diarrhea, nausea and vomiting (14, 15). Interestingly, patients with PACS at 6 months showed gut microbiome dysbiosis compared with non-COVID-19 controls and patients without PACS, while the PACS development was not significantly correlated with viral load both in respiratory and stool (16). Therefore, there is growing emphasis on how butyrate maintaining intestinal homeostasis and reducing lung disruption in SARS-CoV-2 infection.

So far, many therapeutic approaches have been developed for COVID-19 including the use of antiviral drugs, monoclonal antibodies, immunomodulators and convalescent plasma (17–21). However, it is still unclear whether the interventions based on gut microbe and metabolites are effective for the prevention and treatment of respiratory viral infection. As mentioned above, butyrate, a major metabolite of gut microbiota and a fuel for colonocytes, can regulate immune response and mediate gut-lung communication. We hypothesized that oral administration of butyrate protects against SARS-CoV-2 infection. Here, we investigated the effects of butyrate on colon mucosal barrier and lung injury in SARS-CoV-2-challenged golden hamsters. The results showed that butyrate can significantly increase the number of goblet cells in the colon. More importantly, supplementation of butyrate boosted the antiviral immune responses and promoted cell survival in the lung, as a result, alleviated lung injury of SARS-CoV-2-infected hamsters.

## RESULTS

### Body weight change and viral load in hamsters.

To assess whether microbial metabolites protect against virus-induced inflammation and tissue injury, golden hamsters were orally administrated with butyrate before and during the course of SARS-CoV-2 infection (Fig. 1A). From 2 to 5 days post inoculation (dpi), hamsters in the virus-inoculated groups, either butyrate-treated or untreated, showed significant decrease in body weigt, while the mock-infected group showed slight body gain. There was no significant difference between the control and the butyrate-treated groups in weight loss (Fig. 1B). To further investigate the effect of butyrate on SARS-CoV-2 replication, we assessed the viral load in trachea, lung and colon of the hamsters. The highest viral RNA load was detected in the lungs of virus-inoculated individuals throughout the infection (Fig. 1C). The mock infected group had no virus infection (data not shown). From 3 to 5 dpi, viral RNA in the lung was trending higher both in control and butyrate-treated hamsters in contrast to a gradual decrease in the trachea (Fig. 1C). At 5 dpi, viral RNA in the lung of butyrate-treated hamsters was slightly higher than that in the control (P > 0.05) as well as that in the trachea (P > 0.05) (Fig. 1C). Low copies of viral RNA were detected in the colon at 3 and 5 dpi (Fig. 1C). No statistical significance was observed in the viral load between butyrate-treated hamsters and the control (P > 0.05).

**FIG 1.**
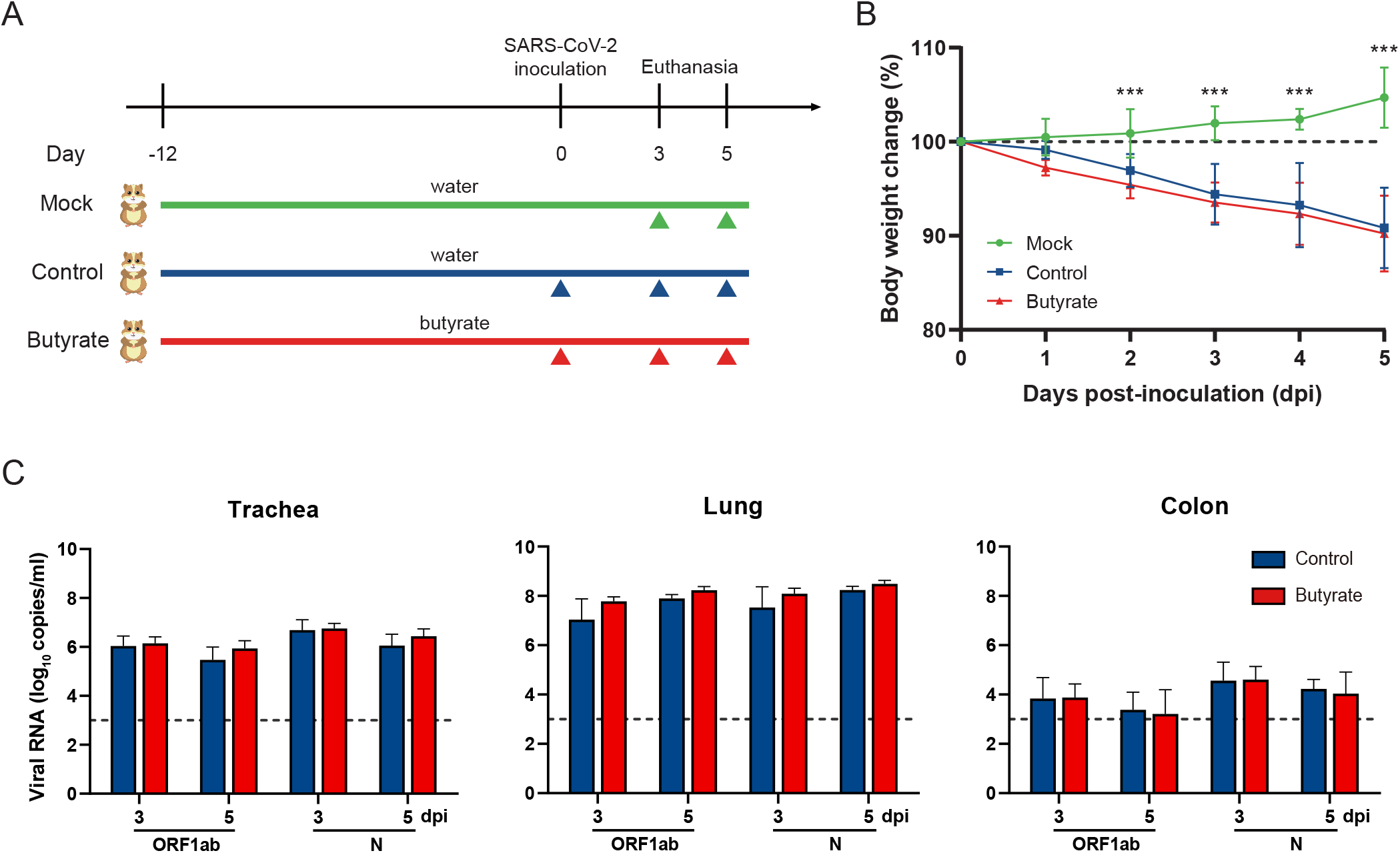
Body weight change and viral load in golden hamsters intranasally challenged with SARS-CoV-2. (A) study design. Hamsters were supplied with or without butyrate in the drinking water since day 12 prior to the virus inoculation till the endpoint of the experiment. Mock animals were hamsters which received pure drinking water and no virus inoculation with SARS-CoV-2. Control indicated hamsters which received drinking water and intranasal inoculation of 1×10^4^ plaque forming units (PFU) of SARS-CoV-2. Butyrate indicated hamsters receiving 500 mmol/L of sodium butyrate supplemented in their daily drinking water and SARS-CoV-2 inoculation. At days 3 (n=3) and 5 (n=8) post-inoculation (dpi), hamsters were euthanized and samples were collected for further analysis. (B) Body weight change after virus inoculation. (C) Viral RNA detected in the trachea, lung and colon of hamsters at 3 and 5 dpi. Data are represented as mean ± SD. Statistical significance were analyzed with two-way ANOVA. *P<0.05, **P<0.01, ***P<0.001. The body weight change and viral load had no significant difference between butyrate-treated hamsters and the control.

### Pathological changes in hamsters.

At 5 dpi, gross observation showed pulmonary hemorrhage and edema in 50-75% in control hamsters, while 10-50% in butyrate-treated hamsters (Fig. 2A). Histopathological analysis of the lung revealed that hamsters treated with butyrate had significantly lower pathological scores with fewer inflammatory cells infiltration, reduced alveolar structure damage and less hemorrhage at 5 dpi (Fig. 2B and C). immunohistochemistry (IHC) for SARS-CoV-2 N Protein (NP) detection showed that there was fewer viral antigen in the lung of butyrate-treated hamsters in comparison to the control (Fig. 2B). As above mentioned, butyrate protected against SARS-CoV-2 by eliminating viral antigen and reducing tissue destruction.

**FIG 2.**
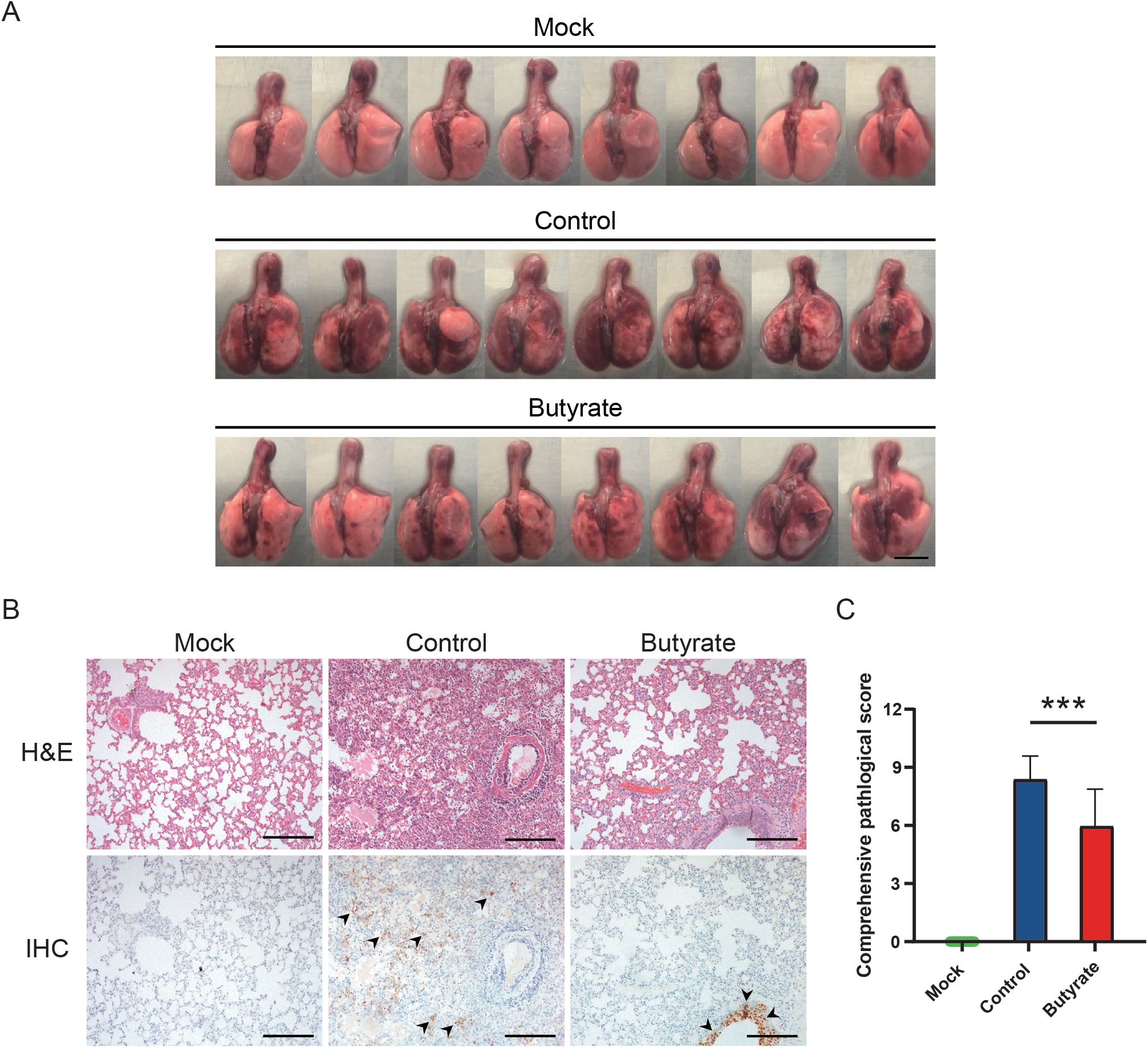
Pathological changes in the lung of golden hamsters intranasally inoculated with SARS-CoV-2. (A) Gross lung images of hamsters at 5 dpi. Scale bars, 1 cm. (B) Histopathological examination of the lungs at 5 dpi. Detection of SARS-CoV-2 NP-positive cells are indicated by black arrows. Scale bars, 200 μm. (C) Comprehensive pathological scores of the lungs at 5 dpi. Data are represented as mean ± SD. Statistical significance were analyzed with Student’s t test. *P<0.05, **P<0.01, ***P<0.001.

### Expression levels of representative genes in hamsters.

To elucidate how butyrate influences antiviral innate immunity, we assessed the expression levels of genes involved following SARS-CoV-2 infection (see Table 1 for primer sequences). First, a marked reduction in interferon alpha and beta receptor subunit 1 gene (*Ifnar1*) was observed in the control compared with both mock and butyrate-treated hamsters at 5 dpi, indicating an inhibited type I interferon (IFN) signaling induced by SARS-CoV-2 (Fig. 3A). Inflammatory cytokines such as interleukin 6 (*Il6*), *Il1b* and tumor necrosis factor alpha (*Tnfα*) were both upregulated in two virus-inoculated groups with higher levels in butyrate-treated hamsters (Fig. 3B). So did the proinflammatory interefone gamma (*Ifng*) (Fig. 3D). Next, to test whether butyrate can influence the immunomodulatory functions of endothelial cells, we assessed mRNA expression of cellular adhesion molecules. Intercellular adhesion molecule 1 (*Icam1*) was marginally downregulated in the control compared with the mock, whereas an increased expression was observed in butyrate-treated hamsters (Fig. 3C). Vascular cell adhesion molecule 1 (*Vcam1*) and selectin E (*Sele*) were both upregulated after SARS-CoV-2 stimulation, but no significant upregulation were seen in *Vcam1* between mock and control hamsters (Fig. 3C). Determination of these adhesion molecules showed that endothelial cells in the lung of butyrate-treated hamsters were activated at 5 dpi. Finally, we found endothelial nitric oxide synthase (*eNOS*, *Nos3*) as well as inducible NOS (*iNOS*, *Nos2*) was significantly decreased in the control compared with the mock, indicating deficient nitric oxide (NO) inside blood vessels upon infection, which thus leads to endothelial dysfunction and suppressed NO signaling in regulating inflammation (Fig. 3D). However, butyrate reversed the downregulation of these two nitric oxide synthases in the lung at 5 dpi (Fig. 3D). Taken together, these results showed that butyrate regulated inflammation by activating antiviral response and promoting homeostasis and activation of endothelial cells.

**FIG 3.**
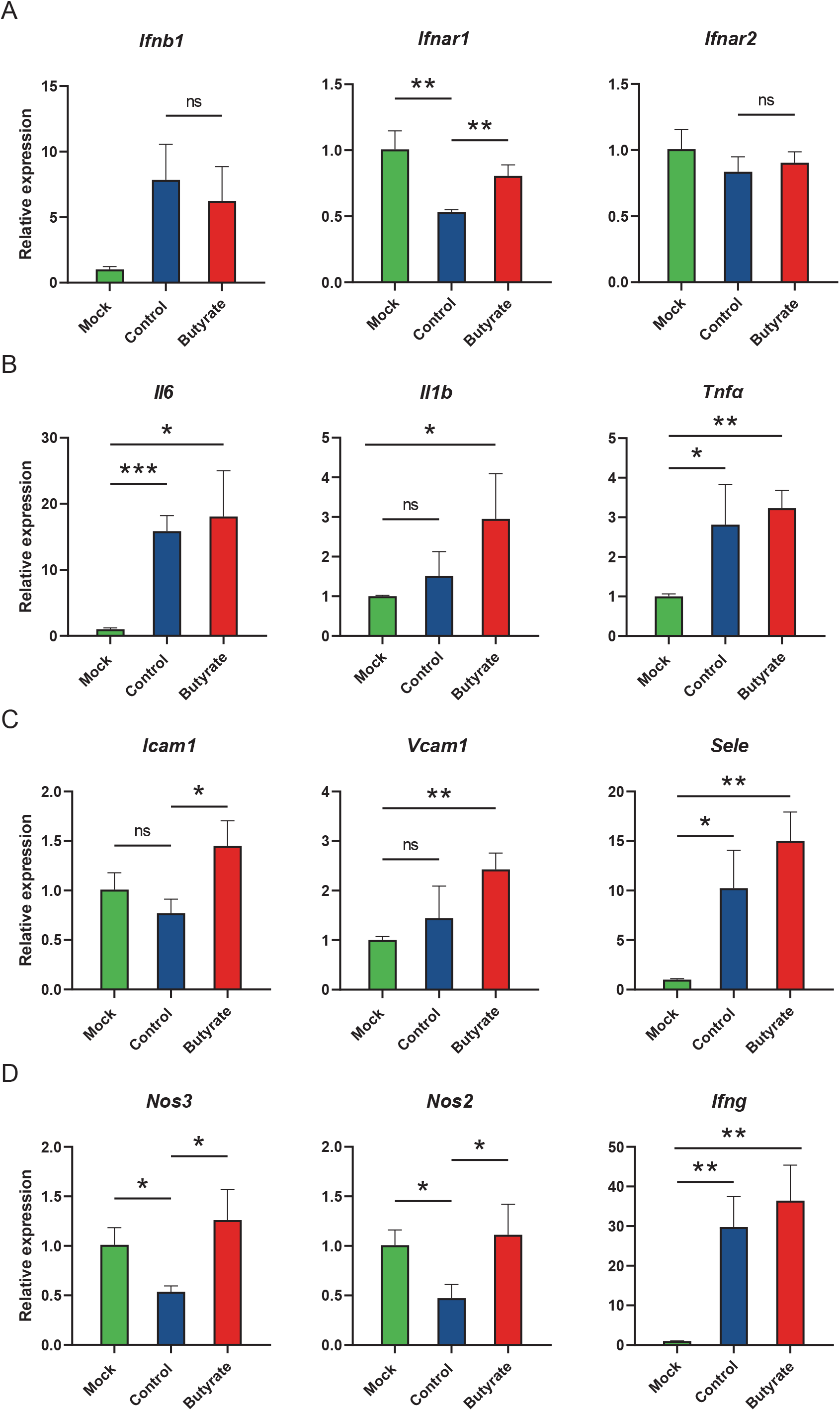
Expression levels of representative genes in golden hamsters intranasally challenged with SARS-CoV-2. Relative mRNA expression for representative genes in (A) type I interferon (IFN) signaling, (B) proinflammatory effect, (C) endothelial cells activation and (D) nitric oxide production in the lungs at 5 dpi. The mRNA level was normalized to the house keeping gene γ-actin and calculated using 2^-ΔΔCt^ method. Data are represented as mean ± SD. Statistical significance was analyzed with Student’s t test. *P< 0.05, **P<0.01, ***P<0.001.

**TABLE 1.**
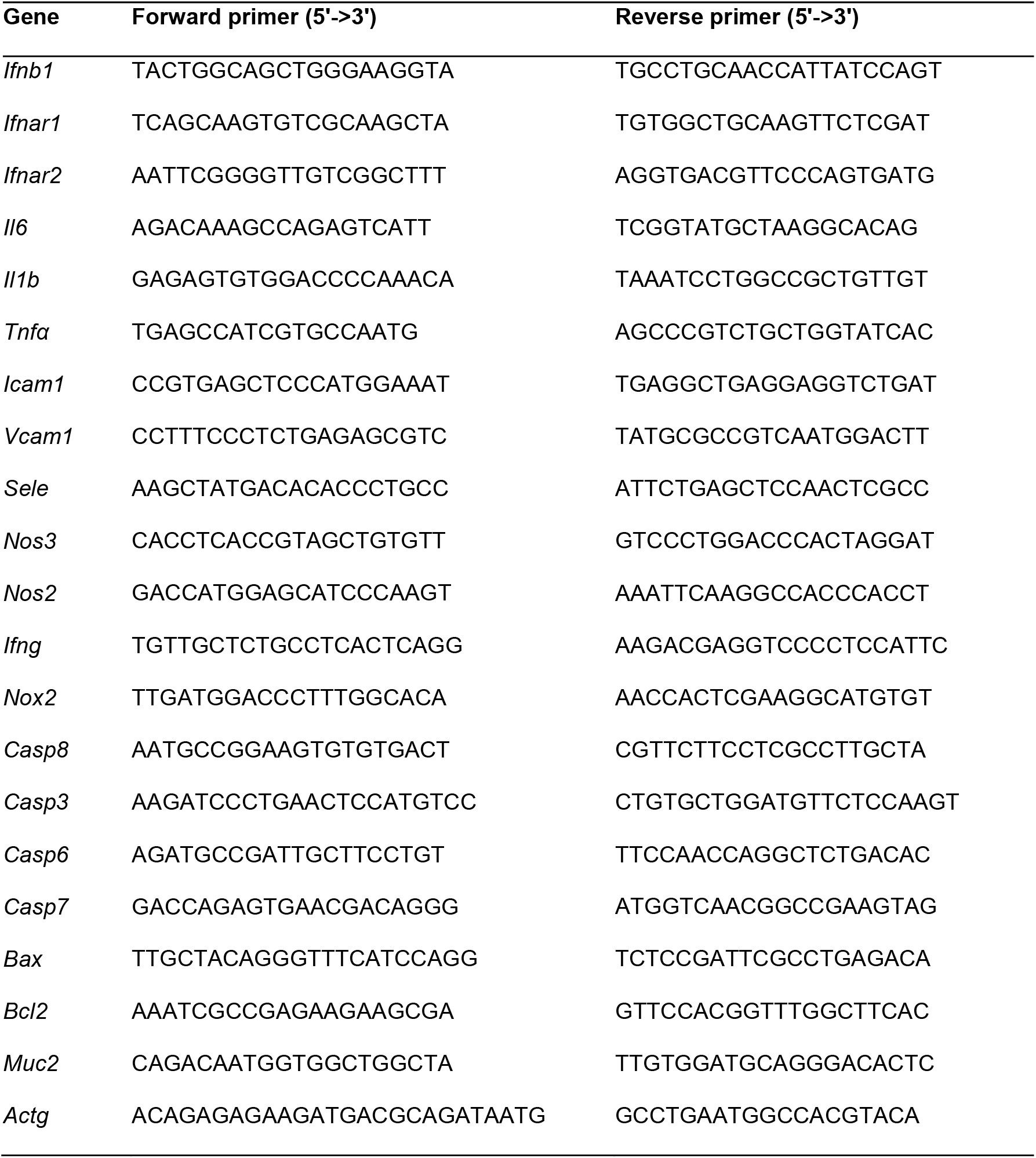
Primer sequences used for RT-qPCR.

### Oxidative status in hamsters.

To further determine the pathogenesis of inflammation, we assessed oxidative stress at 5 dpi. The expression level of NADPH oxidase 2 (*Nox2*) showed slight increase in the control and butyrate-treated hamsters, which probably pointed toward the production of reactive oxygen species (ROS) (Fig. 4A). Compared with the mock, the level of malondialdehyde (MDA) in the plasma of the control was markedly elevated, indicating lipid peroxidation subsequent to oxidative stress (Fig. 4B). Moreover, the activity of superoxide dismutase (SOD), a key antioxidant enzyme in redox signaling, was significantly decreased in the control compared with butyrate-treated hamsters (Fig. 4C). Therefore, butyrate contributed to anti-inflammatory effects through reduction of oxidative stress mainly by regulating redox signaling.

**FIG 4.**
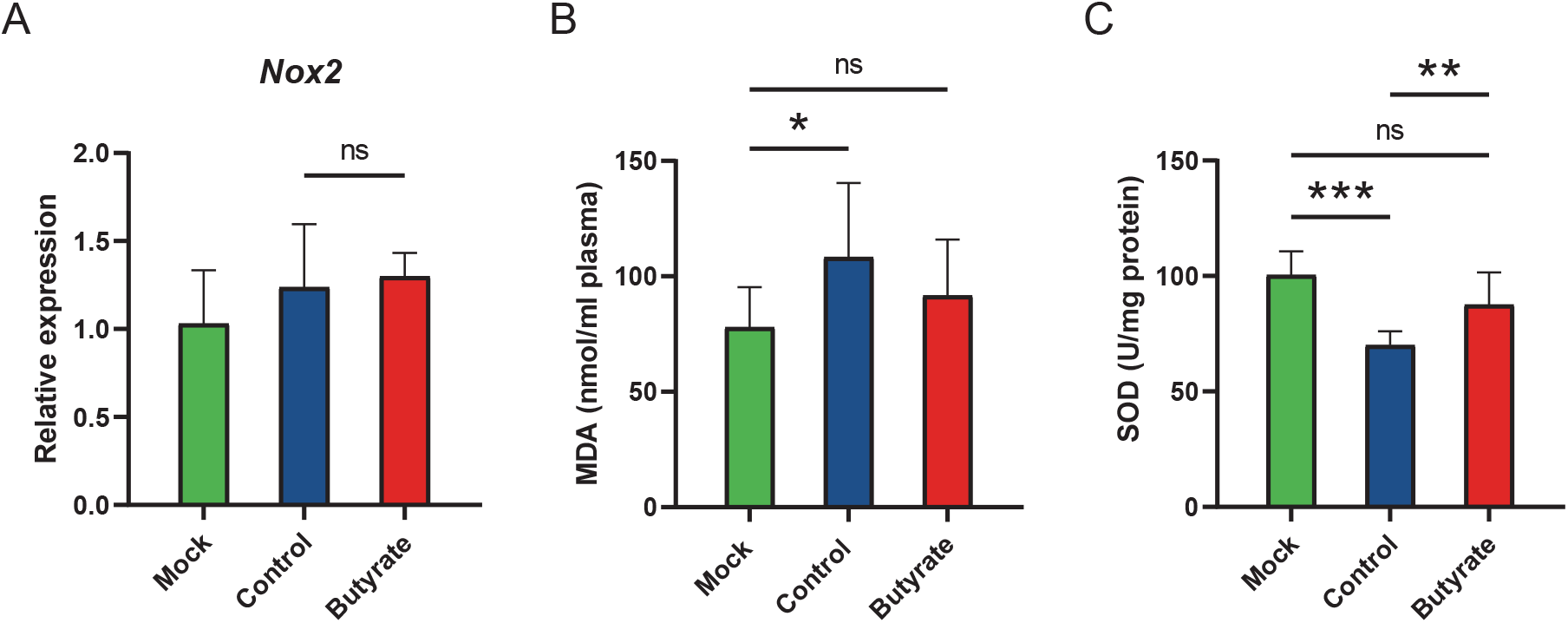
Oxidative status in golden hamsters intranasally challenged with SARS-CoV-2. (A) Relative mRNA expression for NADPH oxidase 2 (*Nox2*) in the lungs at 5 dpi. (B) Malondialdehyde (MDA) levels in the plasma at 5 dpi. (C) Superoxide dismutase (SOD) activity in the lungs at 5 dpi. Data are represented as mean ± SD. Statistical significance was analyzed with Student’s t test. *P<0.05, **P<0.01, ***P<0.001.

### Apoptosis in hamsters.

To determine the consequences of oxidative stress and whether structural integrity of the lung was also affected by butyrate, we assessed SARS-CoV-2-induced apoptosis at 5 dpi. Hoechst staining showed that the lung of butyrate-treated hamsters compared with the control had significantly fewer apoptotic cells (Fig. 5A and B). In line with this, caspase-8 (*Casp8*), a crucial initiator in apoptotic pathway, was significantly increased in the control, but no significant difference between mock and butyrate-treated hamsters (Fig. 5C). Executioner caspase, especially *Casp3*, was upregulated in virus-inoculated groups either butyrate-treated or untreated, indicating apoptosis upon SARS-CoV-2 infection (Fig. 5C). However, the expression of *Bcl2*, an antiapoptotic signature gene, showed significant increase when hamsters were treated with butyrate (Fig. 5C). Thus, butyrate alleviated lung injury by preventing excessive apoptotic cell death and promoting cell survival mediated by an antiapoptotic gene.

**FIG 5.**
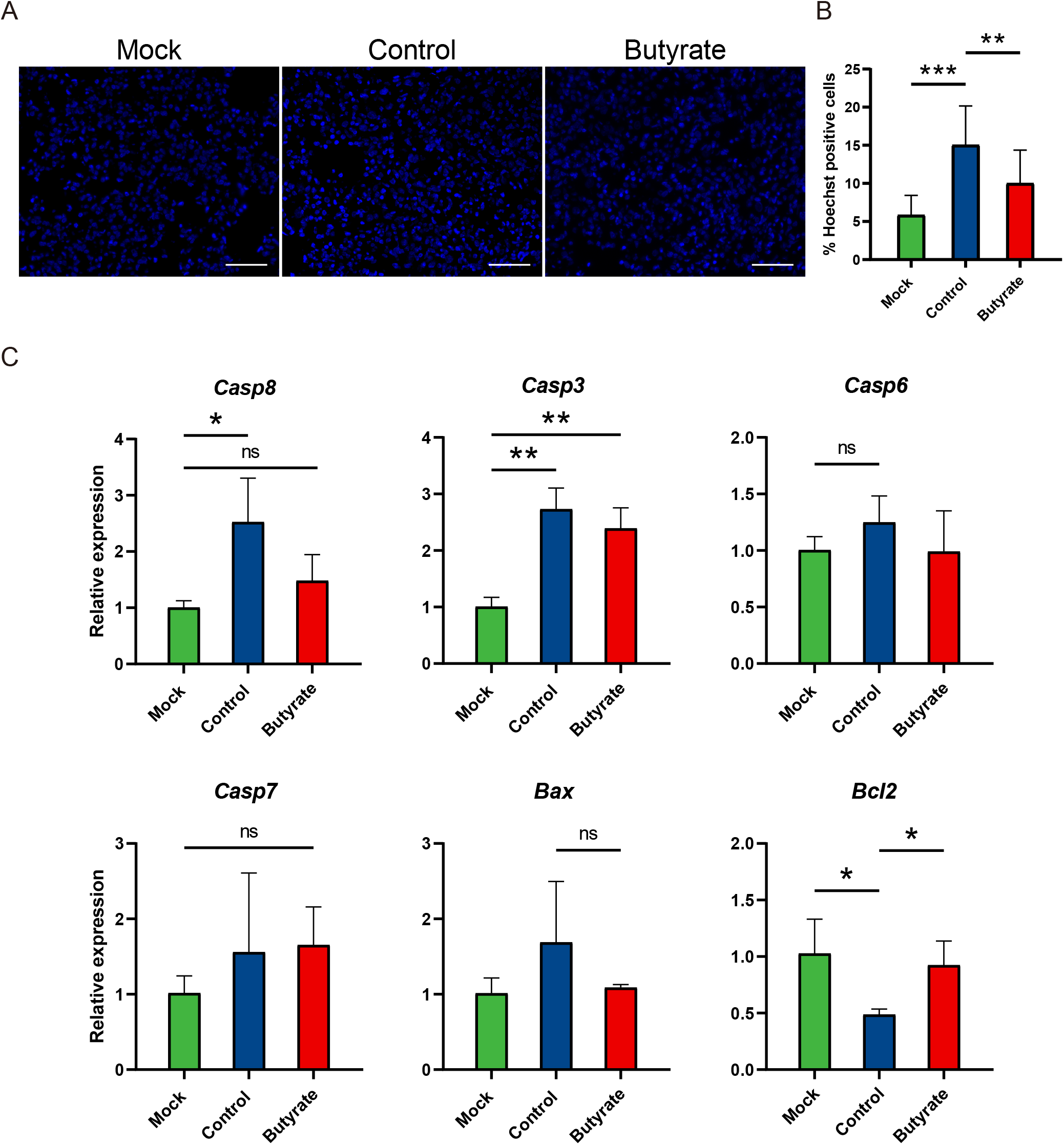
Apoptosis in golden hamsters intranasally challenged with SARS-CoV-2. (A) Hoechst staining of the lungs at 5 dpi. Scale bars, 50 μm. (B) Quantification of Hoechst positive cells in the lungs. (C) Relative mRNA expression for representative genes in apoptosis pathways in the lungs at 5 dpi. The mRNA level was normalized to the house keeping gene γ-actin and calculated using 2^-ΔΔCt^ method. Data are represented as mean ± SD. Statistical significance was analyzed with Student’s t test. *P<0.05, **P<0.01, ***P< 0.001.

### **Goblet cells and *Muc2* expression in hamsters.**

As butyrate is the metabolite mainly produced in colon, we also assessed whether butyrate regulated the development of mucosal barrier. Compared with butyrate-treated hamsters, there were significantly fewer goblet cells in the colon of the control (Fig. 6A and B). Similarly, crypts were elongated (P<0.001) and mucin 2 (*Muc2*) expression was somewhat increased (P>0.05) in butyrate-treated hamsters (Fig. 6B and C). Thus, SARS-CoV-2 infection impaired the colon mucosal barrier and butyrate played a role in goblet cell development.

**FIG 6.**
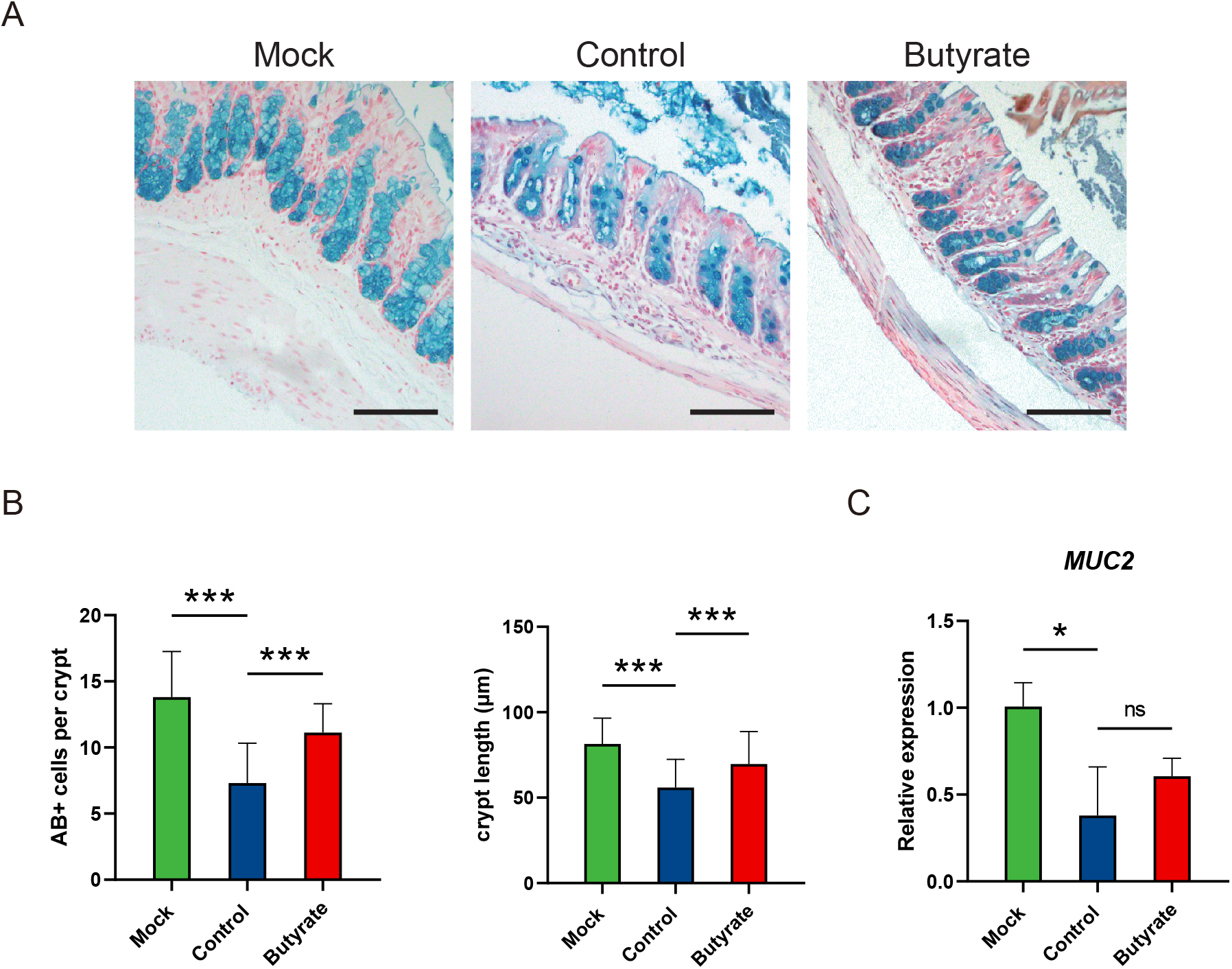
Goblet cells and *Muc2* expression in the colon of golden hamsters intranasally challenged with SARS-CoV-2. (A) AB-stained sections of colon. (B) Quantification of AB+ cells and crypt length in the colon at 5 dpi. Scale bars, 100 μm. (C) *Muc2* expression in the colon at 5 dpi. Data are represented as mean ± SD. Statistical significance was analyzed with Student’s t test. *P<0.05, **P<0.01, ***P<0.001.

## DISCUSSION

Here, we demonstrated that butyrate could protect the SARS-CoV-2-infected hamsters by enhancing antiviral response and promoting cell survival to maintain tissue homeostasis. In severe and critical COVID-19 patients, low or no type I IFNs levels were observed, suggesting a highly impaired type I IFN response in these patients (22, 23). Similarly, the expression of IFNB1 and IFN-stimulated genes (ISGs) such as MX1, ISG20 and OASL failed to be activated in SARS-CoV-2-infected golden hamsters and ferrets respectively (23, 24). At the same time, IFN-I/II receptors-double-knockout mice have increased viral titers and higher congestion scores of the lungs following SARS-CoV-2 infection (25). We found that butyrate activated innate immune response in the early phase of SARS-CoV-2 infection, characterized by upregulated type I IFN signaling and increased proinflammatory cytokines, which contributed to rapid viral antigen clearance and avoided immunopathology in the lungs. This was also supported by a recent study which showed that antiviral factors such as IL1b, IRF7, TNF and IFNAR1 were upregulated in butyrate-treated gut epithelial organoids (26).

Another feature of severe COVID-19 patients was lymphopenia and immunosuppression, which was responsible for hyperinflammation in the late stage of disease (27, 28). As innate immune response alone may be insufficient to viral clearance, recruitment of lymphocytes appeared a more effective defense in virus infection. Alveolar capillary endothelial cells not only functioned as gas change, but were capable of recruiting immune cells through adhesion molecules and activating CD4+ T cells (29). By binding to T cell integrin, ICAM1 increased T cell receptor (TCR) signaling to mediate activation, adhesion and migration of T cells (30, 31). Patients with COVID-19 showed pulmonary vascular injury associated with intracellular presence of SARS-CoV-2 and endothelial cell destruction (32). In a SARS-CoV-2-infected vascularized lung-on-chip model, despite unproductive viral replication, lower CD31 expression and decreased barrier integrity were observed (33). These evidences indicated endothelial injury might lead to impaired immune cell recruitment and thus increased hyperinflammation in the lung. In our study, SARS-CoV-2-infected hamsters had decreased gene expression levels of adhesion molecules (*Icam1* and *Vcam1*), suggesting suppression of endothelial cells activation. NOS3 was mainly expressed in endothelial cells and NOS3-derived NO was involved in maintaining vascular homeostasis, like vasodilation, inhibition of vascular inflammation and preventing endothelial cells apoptosis (34). Together with NOS2, endogenous NO produced also regulated T cell differentiation and activation (35). Our data showed that butyrate reversed the expression of *Nos3* and *Nos2* induced by SARS-CoV-2. Thus, butyrate offered endothelial protection and promoted endothelial cells activation.

Further exploring the effect of butyrate on reducing tissue damage, we observed that butyrate played anti-oxidative and anti-apoptotic effects. Increasing evidences suggested that pathological responses in COVID-19 patients was probably caused by oxidative stress (36). Excessive ROS and subsequent MDA, a lipid peroxidation product, were both oxidative markers (37, 38). Moreover, there is decreased expression of the antioxidant enzyme SOD3 in the lungs of elderly COVID-19 patients (39). Not only that, the link between oxidative stress and apoptosis has been proven (40). Apoptosis induced by SARS-CoV-2 was associated with disease severity and inhibition of intrinsic apoptosis could markedly ameliorated the lung damage in transgenic mice that expressed human angiotensin-converting enzyme 2 (hACE2) (41, 42). BCL2 is known to suppress apoptosis by regulating ROS levels in cytoplasm and mitochondria (40). Thus, our findings suggested that butyrate might inhibit SARS-CoV-2-induced apoptosis by improving antioxidant capacity in the lung.

In summary, we demonstrated that butyrate protected against SARS-CoV-2-induced tissue damage in golden hamsters. Among respiratory diseases, butyrate has previously been associated with regulation in chronic pulmonary disorders and no significant effects are observed in treating SARS-CoV-2-infected hamsters with a combination of SCFAs (i.e. acetate, propionate and butyrate) (43, 44). Our study highlighted the beneficial effects of butyrate on boosting antiviral immune response and reducing oxidative stress to promote cell survival in the disease.

## MATERIALS AND METHODS

### Virus

The SARS-CoV-2 D614G variant AP62 (hCoV-19/China/AP62/2020, GISAID accession No. EPI_ISL_2779638) was used in this study. Virus stocks were prepared by three passages in Vero (ATCC CCL-81) in Dulbecco’s modified Eagle Medium (DMEM) (Gibco) with 1% Penicillin-Streptomycin (Gibco). Virus titers were measured by plaque assay.

### Experimental animal and study design

8-10-week-old male golden hamsters were derived from Charles River Laboratories (Beijing Vital River Laboratory Animal Technology Co., Ltd.) and raised at the specific pathogen-free animal feeding facilities. For butyrate-treated group, sodium butyrate (Sigma-Aldrich) was supplemented in the drinking water at a final concentration of 500 mmol/L 12 days prior to virus inoculation and until the end of the experiment (3 or 5 days post-inoculation, dpi) (Figure 1). Control hamsters were supplied with water without butyrate during the experiment. Hamsters were anaesthetized with isoflurane and intranasally inoculated with a dose of 1×10^4^ plaque forming units (PFU) of SARS-CoV-2 diluted in 200 μL phosphate-buffered saline (PBS). Mock animals were inoculated with 200 μL PBS. Body weight of each hamster was measured daily during the course of the experiment. At day 3 and 5 post-inoculation, three and eight hamsters were euthanized respectively. Blood samples were collected to prepare plasma. After gross observation and pathological examination, trachea, lung and colon were collected to determine viral load or levels of host gene expression. Lung and colon tissues were also fixed in 10% formalin for histologic analysis. All experiments with the infectious virus were performed in biosafety level 3 (BSL-3) and animal biosafety level 3 (ABSL-3) containment facilities. The animal experiment was approved by the Medical Animal Care and Welfare Committee of Shantou University Medical College (Ref No. SUMC2023-058).

### Determination of viral load

Fresh trachea, lung and colon tissues were collected and homogenized in PBS (100 mg/mL) respectively and RNA was extracted using the RNA Extraction Kit (Wantai Beijing). Quantitative real-time PCR (RT-qPCR) was performed to detect the ORF1ab and N gene of SARS-CoV-2 using the SARS-CoV-2 RT-qPCR Kit (Wantai Beijing) on a SLAN-96S real-time PCR system (Hongshi Shanghai).

### Determination of host gene expression level

Tissues kept in Invitrogen^TM^ RNAlater^TM^ Stabilization Solution (Thermo Fisher Scientific) were homogenized in buffer RLT Plus (Qiagen). Total RNA from lung and colon samples was extracted using RNeasy Plus Mini Kit (Qiagen) according to the manufacturer’s instructions. For determination of target gene expression level, cDNA was synthetized from total RNA using PrimeScript II 1st Strand cDNA Synthesis Kit (Takara) and amplified using ChamQ Universal SYBR qPCR Master Mix (Vazyme Biotech) on a SLAN-96S real-time PCR system (Hongshi Shanghai). Primer sequences used for amplification were listed in Table 1. For each sample, the host gene expression level was normalized to the house keeping gene γ-actin (*Actg*) and calculated using 2^-ΔΔCt^ method.

### Histologic analysis

After being fixed in 10% formalin, lung and colon tissues were then embedded into paraffin and sectioned into 3-5 μm slices. The fixed lung sections were stained with hematoxylin and eosin (H&E) for histopathological analysis. Lung injury was evaluated according to pathological changes as follows: 1) alveolar septum widened and consolidation; 2) pulmonary hemorrhage and edema; and 3) inflammatory cell infiltration. Score each pathological change on a scale of 0 to 4: 0=no damage; 1=mild injury; 2 =moderate injury; 3=severe injury; and 4=very severe (45). For one lung lobe, the pathological score was the sum of these pathological changes. For each hamster, the comprehensive pathological score was averaged over the pathological score of 3 or 4 lung lobes. To elucidate the distribution of viral antigen in lung tissues, immunohistochemistry (IHC) was used to detect the N protein (NP) of SARS-CoV-2. A murine anti-SARS-CoV-2 NP specific monoclonal antibody (15A7-1) was applied (45). The colon sections were stained with Alcian Blue (AB) to analyze goblet cells and crypt length. For each hamster, the number of goblet cells and the crypt length were averaged over at least 5 well-defined crypts (46). Images were taken using a ZEISS Axio Imager A2 microscope.

### Apoptosis assay

Formalin-fixed lung sections were stained with Hoechst 33258 (Beyotime) and the images were captured using a fluorescence microscope (ZEISS, Axio Imager A2). Apoptotic cells are characterized by condensed chromatin, so the cells which showed higher fluorescence intensity in nuclei were considered as Hoechst positive cells. The percentage of Hoechst positive cells was measured in ImageJ (NIH).

### Malondialdehyde (MDA) and superoxide dismutase (SOD) assays

Plasma prepared was used for MDA measurement with Lipid Peroxidation MDA Assay Kit (Beyotime). Freshly collected lung samples were homogenized in lysis buffer (Beyotime) for SOD detection by Superoxide Dismutase (SOD) Assay Kit (Nanjing Jiancheng). One unit of SOD is defined as the amount of enzyme that causes 50% inhibition of the reduction reaction between water-soluble tetrazolium salt-1 (WST-1) and superoxide anion.

### Statistics

All results were presented as mean ± standard deviation (SD). Student’s t test (two tailed) and two-way analysis of variance (ANOVA) was used for comparison of treatment groups. Mann-Whitney test was used to calculate the P value in the case of non-normal distribution of the data. P<0.05 was statistically significant. *P < 0.05, **P < 0.01, *** P < 0.001. NS = no significance. Graph generation and statistical analysis were performed in GraphPad Prism 8.0.1 (GraphPad Software).

## ACKNOWLEDGMENTS

We thank all staff from the Guangdong-Hong Kong Joint Laboratory of Emerging Infectious Diseases / Joint Laboratory for International Collaboration in Virology and Emerging Infectious Diseases / Joint Institute of Virology (STU/HKU) and SKLEID for their technical support and administrative assistance.

This research was funded by Shenzhen-Hong Kong Science and Technology Cooperation Zone-Shenzhen program (grant number HZQB-KCZYZ-2021014), Department of Science & Technology, Guangdong (grant number 2019B121205009), Hong Kong Research Grant Council (grant number T11-705/21-N and T11-712/19-N), the Innovation and Technology Commission of Hong Kong and Li Ka Shing Foundation. The funders had no role in the study design, data collection and analysis, decision to publish, or preparation of the article.

The authors declare no competing interests. Conceptualization, H.Y. and H.Z.; methodology, H.Y., L.Y. and H.Z.; resources and supervision, L.Y., N.X., Y.G. and H.Z.; experimental investigation, H.Y., L.Y., Z.Y., M.Z., J.Y., K.W., W.C. and R.C.; formal analysis, H.Y. and H.Z.; visualization, H.Y.; manuscript writing, H.Y. (original draft) and H.Z.; project administration, H.Z.; funding acquisition, Y.G. and H.Z..

## REFERENCES

1. Zhang L, Liu C, Jiang Q, Yin Y. 2021. Butyrate in Energy Metabolism: There Is Still More to Learn. Trends Endocrinol Metab 32:159–169.

2. Medawar E, Haange SB, Rolle-Kampczyk U, Engelmann B, Dietrich A, Thieleking R, Wiegank C, Fries C, Horstmann A, Villringer A, von Bergen M, Fenske W, Veronica Witte A. 2021. Gut microbiota link dietary fiber intake and short-chain fatty acid metabolism with eating behavior. Transl Psychiatry 11:500.

3. Koh A, De Vadder F, Kovatcheva-Datchary P, Bäckhed F. 2016. From Dietary Fiber to Host Physiology: Short-Chain Fatty Acids as Key Bacterial Metabolites. Cell 165:1332–1345.

4. Chang PV, Hao L, Offermanns S, Medzhitov R. 2014. The microbial metabolite butyrate regulates intestinal macrophage function via histone deacetylase inhibition. Proc Natl Acad Sci U S A 111:2247–52.

5. Furusawa Y, Obata Y, Fukuda S, Endo TA, Nakato G, Takahashi D, Nakanishi Y, Uetake C, Kato K, Kato T, Takahashi M, Fukuda NN, Murakami S, Miyauchi E, Hino S, Atarashi K, Onawa S, Fujimura Y, Lockett T, Clarke JM, Topping DL, Tomita M, Hori S, Ohara O, Morita T, Koseki H, Kikuchi J, Honda K, Hase K, Ohno H. 2013. Commensal microbe-derived butyrate induces the differentiation of colonic regulatory T cells. Nature 504:446–50.

6. Wei H, Yu C, Zhang C, Ren Y, Guo L, Wang T, Chen F, Li Y, Zhang X, Wang H, Liu J. 2023. Butyrate ameliorates chronic alcoholic central nervous damage by suppressing microglia-mediated neuroinflammation and modulating the microbiome-gut-brain axis. Biomed Pharmacother 160:114308.

7. Dang AT, Marsland BJ. 2019. Microbes, metabolites, and the gut-lung axis. Mucosal Immunol 12:843–850.

8. Tilg H, Adolph TE, Trauner M. 2022. Gut-liver axis: Pathophysiological concepts and clinical implications. Cell Metab 34:1700–1718.

9. WHO. 2020. The top 10 causes of death. https://www.who.int/news-room/fact-sheets/detail/the-top-10-causes-of-death.

10. WHO. 2023. WHO Coronavirus (COVID-19) Dashboard. https://covid19.who.int.

11. Wolfel R, Corman VM, Guggemos W, Seilmaier M, Zange S, Muller MA, Niemeyer D, Jones TC, Vollmar P, Rothe C, Hoelscher M, Bleicker T, Brunink S, Schneider J, Ehmann R, Zwirglmaier K, Drosten C, Wendtner C. 2020. Virological assessment of hospitalized patients with COVID-2019. Nature 581:465–469.

12. V’Kovski P, Kratzel A, Steiner S, Stalder H, Thiel V. 2021. Coronavirus biology and replication: implications for SARS-CoV-2. Nature reviews Microbiology 19:155–170.

13. Lamers MM, Haagmans BL. 2022. SARS-CoV-2 pathogenesis. Nat Rev Microbiol 20:270–284.

14. Patel KP, Patel PA, Vunnam RR, Hewlett AT, Jain R, Jing R, Vunnam SR. 2020. Gastrointestinal, hepatobiliary, and pancreatic manifestations of COVID-19. J Clin Virol 128:104386.

15. Cooney J, Poullis A. 2022. Post-COVID-19 irritable bowel syndrome. Neurogastroenterol Motil 34:e14420.

16. Liu Q, Mak JWY, Su Q, Yeoh YK, Lui GC-Y, Ng SSS, Zhang F, Li AYL, Lu W, Hui DS-C, Chan PK, Chan FKL, Ng SC. 2022. Gut microbiota dynamics in a prospective cohort of patients with post-acute COVID-19 syndrome. Gut 71:544–552.

17. Saravolatz LD, Depcinski S, Sharma M. 2023. Molnupiravir and Nirmatrelvir-Ritonavir: Oral Coronavirus Disease 2019 Antiviral Drugs. Clin Infect Dis 76:165–171.

18. Taylor PC, Adams AC, Hufford MM, de la Torre I, Winthrop K, Gottlieb RL. 2021. Neutralizing monoclonal antibodies for treatment of COVID-19. Nat Rev Immunol 21:382–393.

19. Korley FK, Durkalski-Mauldin V, Yeatts SD, Schulman K, Davenport RD, Dumont LJ, El Kassar N, Foster LD, Hah JM, Jaiswal S, Kaplan A, Lowell E, McDyer JF, Quinn J, Triulzi DJ, Van Huysen C, Stevenson VLW, Yadav K, Jones CW, Kea B, Burnett A, Reynolds JC, Greineder CF, Haas NL, Beiser DG, Silbergleit R, Barsan W, Callaway CW, Investigators S-CP. 2021. Early Convalescent Plasma for High-Risk Outpatients with Covid-19. N Engl J Med 385:1951–1960.

20. Rossotti R, Travi G, Ughi N, Corradin M, Baiguera C, Fumagalli R, Bottiroli M, Mondino M, Merli M, Bellone A, Basile A, Ruggeri R, Colombo F, Moreno M, Pastori S, Perno CF, Tarsia P, Epis OM, Puoti M, Niguarda C-WG. 2020. Safety and efficacy of anti-il6-receptor tocilizumab use in severe and critical patients affected by coronavirus disease 2019: A comparative analysis. The Journal of infection doi:10.1016/j.jinf.2020.07.008.

21. Group RC, Horby P, Lim WS, Emberson JR, Mafham M, Bell JL, Linsell L, Staplin N, Brightling C, Ustianowski A, Elmahi E, Prudon B, Green C, Felton T, Chadwick D, Rege K, Fegan C, Chappell LC, Faust SN, Jaki T, Jeffery K, Montgomery A, Rowan K, Juszczak E, Baillie JK, Haynes R, Landray MJ. 2021. Dexamethasone in Hospitalized Patients with Covid-19. N Engl J Med 384:693–704.

22. Hadjadj J, Yatim N, Barnabei L, Corneau A, Boussier J, Smith N, Péré H, Charbit B, Bondet V, Chenevier-Gobeaux C, Breillat P, Carlier N, Gauzit R, Morbieu C, Pène F, Marin N, Roche N, Szwebel T-A, Merkling SH, Treluyer J-M, Veyer D, Mouthon L, Blanc C, Tharaux P-L, Rozenberg F, Fischer A, Duffy D, Rieux-Laucat F, Kernéis S, Terrier B. 2020. Impaired type I interferon activity and inflammatory responses in severe COVID-19 patients. Science (New York, NY) 369:718–724.

23. Blanco-Melo D, Nilsson-Payant BE, Liu WC, Uhl S, Hoagland D, Møller R, Jordan TX, Oishi K, Panis M, Sachs D, Wang TT, Schwartz RE, Lim JK, Albrecht RA, tenOever BR. 2020. Imbalanced Host Response to SARS-CoV-2 Drives Development of COVID-19. Cell 181:1036–1045.e9.

24. Hoagland DA, Møller R, Uhl SA, Oishi K, Frere J, Golynker I, Horiuchi S, Panis M, Blanco-Melo D, Sachs D, Arkun K, Lim JK, tenOever BR. 2021. Leveraging the antiviral type I interferon system as a first line of defense against SARS-CoV-2 pathogenicity. Immunity 54.

25. Leist SR, Dinnon KH, 3rd, Schäfer A, Tse LV, Okuda K, Hou YJ, West A, Edwards CE, Sanders W, Fritch EJ, Gully KL, Scobey T, Brown AJ, Sheahan TP, Moorman NJ, Boucher RC, Gralinski LE, Montgomery SA, Baric RS. 2020. A Mouse-Adapted SARS-CoV-2 Induces Acute Lung Injury and Mortality in Standard Laboratory Mice. Cell doi:10.1016/j.cell.2020.09.050.

26. Li J, Richards EM, Handberg EM, Pepine CJ, Raizada MK. 2021. Butyrate Regulates COVID-19-Relevant Genes in Gut Epithelial Organoids From Normotensive Rats. Hypertension (Dallas, Tex : 1979) 77:e13-e16.

27. Yu K, He J, Wu Y, Xie B, Liu X, Wei B, Zhou H, Lin B, Zuo Z, Wen W, Xu W, Zou B, Wei L, Huang X, Zhou P. 2020. Dysregulated adaptive immune response contributes to severe COVID-19. Cell Res 30:814–816.

28. Tian W, Zhang N, Jin R, Feng Y, Wang S, Gao S, Gao R, Wu G, Tian D, Tan W, Chen Y, Gao GF, Wong CCL. 2020. Immune suppression in the early stage of COVID-19 disease. Nat Commun 11:5859.

29. Gillich A, Zhang F, Farmer CG, Travaglini KJ, Tan SY, Gu M, Zhou B, Feinstein JA, Krasnow MA, Metzger RJ. 2020. Capillary cell-type specialization in the alveolus. Nature 586:785–789.

30. Jankowska KI, Williamson EK, Roy NH, Blumenthal D, Chandra V, Baumgart T, Burkhardt JK. 2018. Integrins Modulate T Cell Receptor Signaling by Constraining Actin Flow at the Immunological Synapse. Front Immunol 9:25.

31. Johansen KH, Golec DP, Huang B, Park C, Thomsen JH, Preite S, Cannons JL, Garcon F, Schrom EC, Courreges CJF, Veres TZ, Harrison J, Nus M, Phelan JD, Bergmeier W, Kehrl JH, Okkenhaug K, Schwartzberg PL. 2022. A CRISPR screen targeting PI3K effectors identifies RASA3 as a negative regulator of LFA-1-mediated adhesion in T cells. Sci Signal 15:eabl9169.

32. Capuano A, Rossi F, Paolisso G. 2020. Covid-19 Kills More Men Than Women: An Overview of Possible Reasons. Front Cardiovasc Med 7:131.

33. Thacker VV, Sharma K, Dhar N, Mancini GF, Sordet-Dessimoz J, McKinney JD. 2021. Rapid endotheliitis and vascular damage characterize SARS-CoV-2 infection in a human lung-on-chip model. EMBO Rep 22:e52744.

34. Forstermann U, Sessa WC. 2012. Nitric oxide synthases: regulation and function. Eur Heart J 33:829–37, 837a-837d.

35. Garcia-Ortiz A, Serrador JM. 2018. Nitric Oxide Signaling in T Cell-Mediated Immunity. Trends Mol Med 24:412–427.

36. Laforge M, Elbim C, Frere C, Hemadi M, Massaad C, Nuss P, Benoliel JJ, Becker C. 2020. Tissue damage from neutrophil-induced oxidative stress in COVID-19. Nat Rev Immunol 20:515–516.

37. Forman HJ, Zhang H. 2021. Targeting oxidative stress in disease: promise and limitations of antioxidant therapy. Nat Rev Drug Discov 20:689–709.

38. Cen M, Ouyang W, Zhang W, Yang L, Lin X, Dai M, Hu H, Tang H, Liu H, Xia J, Xu F. 2021. MitoQ protects against hyperpermeability of endothelium barrier in acute lung injury via a Nrf2-dependent mechanism. Redox Biol 41:101936.

39. Abouhashem AS, Singh K, Azzazy HME, Sen CK. 2020. Is Low Alveolar Type II Cell SOD3 in the Lungs of Elderly Linked to the Observed Severity of COVID-19? Antioxid Redox Signal 33:59–65.

40. Sharma P, Kaushal N, Saleth LR, Ghavami S, Dhingra S, Kaur P. 2023. Oxidative stress-induced apoptosis and autophagy: Balancing the contrary forces in spermatogenesis. Biochim Biophys Acta Mol Basis Dis 1869:166742.

41. Andre S, Picard M, Cezar R, Roux-Dalvai F, Alleaume-Butaux A, Soundaramourty C, Cruz AS, Mendes-Frias A, Gotti C, Leclercq M, Nicolas A, Tauzin A, Carvalho A, Capela C, Pedrosa J, Castro AG, Kundura L, Loubet P, Sotto A, Muller L, Lefrant JY, Roger C, Claret PG, Duvnjak S, Tran TA, Racine G, Zghidi-Abouzid O, Nioche P, Silvestre R, Droit A, Mammano F, Corbeau P, Estaquier J. 2022. T cell apoptosis characterizes severe Covid-19 disease. Cell Death Differ 29:1486–1499.

42. Chu H, Shuai H, Hou Y, Zhang X, Wen L, Huang X, Hu B, Yang D, Wang Y, Yoon C, Wong BH, Li C, Zhao X, Poon VK, Cai JP, Wong KK, Yeung ML, Zhou J, Au-Yeung RK, Yuan S, Jin DY, Kok KH, Perlman S, Chan JF, Yuen KY. 2021. Targeting highly pathogenic coronavirus-induced apoptosis reduces viral pathogenesis and disease severity. Sci Adv 7.

43. Corrêa RO, Castro PR, Moser R, Ferreira CM, Quesniaux VFJ, Vinolo MAR, Ryffel B. 2022. Butyrate: Connecting the gut-lung axis to the management of pulmonary disorders. Front Nutr 9:1011732.

44. Sencio V, Machelart A, Robil C, Benech N, Hoffmann E, Galbert C, Deryuter L, Heumel S, Hantute-Ghesquier A, Flourens A, Brodin P, Infanti F, Richard V, Dubuisson J, Grangette C, Sulpice T, Wolowczuk I, Pinet F, Prévot V, Belouzard S, Briand F, Duterque-Coquillaud M, Sokol H, Trottein F. 2022. Alteration of the gut microbiota following SARS-CoV-2 infection correlates with disease severity in hamsters. Gut microbes 14:2018900.

45. Yuan L, Zhu H, Zhou M, Ma J, Liu X, Wu K, Ye J, Yu H, Chen P, Chen R, Wang J, Zhang Y, Ge S, Yuan Q, Cheng T, Guan Y, Xia N. 2022. Nasal irrigation efficiently attenuates SARS-CoV-2 Omicron infection, transmission and lung injury in the Syrian hamster model. iScience 25:105475.

46. Johansson ME, Hansson GC. 2022. Goblet cells need some stress. J Clin Invest 132.

